# The functional anatomy of self-generated and predictable speech

**DOI:** 10.1101/119644

**Authors:** Lena K. L. Oestreich, Thomas J. Whitford, Marta I. Garrido

## Abstract

Sensory attenuation refers to the cortical suppression of self-generated sensations relative to externally-generated sensations. This attenuation of cortical responsiveness is the result of internal forward models which make precise predictions about forthcoming sensations. Forward models of sensory attenuation in the auditory domain are thought to operate along auditory white matter pathways such as the arcuate fasciculus and the frontal aslant. The aim of this study was to investigate whether brain regions that are structurally connected via these white matter pathways are also effectively connected during overt speech, as well as as when listening to externally-generated speech that is temporally predictable via a visual cue. Using Electroencephalography (EEG) and Dynamic Causal Modelling (DCM) we investigated network models that link the primary auditory cortex to Wernicke’s and Broca’s area either directly or indirectly through Geschwind’s territory, which are structurally connected via the arcuate fasciculus. Connections between Broca’s and supplementary motor area, which are structurally connected via the frontal aslant, were also included. Our results revealed that bilateral areas interconnected by indirect and direct pathways of the arcuate fasciculus, in addition to regions interconnected by the frontal aslant best explain the EEG responses to both self-generated speech, and speech that is externally-generated but temporally predictable. These findings indicate that structurally connected brain regions thought to be involved in auditory attenuation are also effectively connected. Critically, our findings expand on the notion of internal forward models, whereby sensory consequences of our actions are internally predicted and reflected in reduced cortical responsiveness to these sensations.

**Highlights:**

- Effective connectivity of auditory attenuation to self-generated and predictable speech
- EEG and DCM were used to investigate several plausible network models
- Structurally connected brain areas of auditory attenuation are effectively connected
- Internal forward models modulate self-generated and predictable speech

## 1. Introduction

The ability to predict imminent auditory sensations that are either self-generated in the form of speech, or based on past experiences such as hearing a familiar song, is crucial for processing the abundance of auditory stimulation we experience at any moment, as it helps us to adapt to unexpected auditory events in the environment. Sensory attenuation refers to the reduction in the neurophysiological response to sensations that are generated by our own actions relative to identical sensations that are generated in the external environment. (Wolpert et al., 1995) suggested that sensory attenuation is the result of internal forward models, whereby the sensory consequences of our own actions are predicted on the basis of an efference copy of the motor command. Due to the predicted sensory consequences of internally generated actions, the central nervous system tends to be less responsive to self-generated actions as opposed to identical sensations that are externally generated.

Blakemore et al. (1998) postulated that we cannot tickle ourselves because an efference copy of the motor command is sent to the sensory cortex where, as a consequence, the responsiveness to the forthcoming tickle-sensation is suppressed. Likewise, self-generated eye-movements have been demonstrated to elicit an efference copy, which is sent to the visual cortex (Sperry, 1950). This facilitates that rather than perceiving the room to be moving, we recognize that the changes in visual input are due to eye movements(Sperry, 1950). Similarly, speech production areas in the frontal lobes send efference copies of self-generated speech sounds to the auditory cortex (Creutzfeldt et al., 1989), which predict the onset of self-generated overt speech (and arguably also inner speech). The auditory cortical responsiveness is therefore attenuated, which enables the recognition of speech sounds (and arguably also inner speech sounds) to be self-generated (Feinberg, 1978; Ford et al., 2007).

Eliades and Wang (2003) provided *in vivo* evidence for internal forward models in marmosets by directly recording from the auditory cortex using intracranial electrodes. It was reported that activity of the neurons in the auditory cortex was significantly suppressed while the primates engaged in willed vocalizations. These findings are in line with electroencephalography (EEG) reports of auditory cortical attenuation during willed vocalization in humans: Healthy individuals exhibit significantly reduced cortical responsiveness to sounds that are self-versus externally generated (Schafer and Marcus, 1973). This is reflected in a reduced N1 amplitude, an auditory evoked potential component, which peaks at approximately 100ms after the onset of a sound and is reportedly elicited in the auditory cortex (Zouridakis et al., 1998). Auditory attenuation has been observed for willed vocalizations (Curio et al., 2000; Heinks-Maldonado et al., 2005), button-press elicited sounds (Schafer and Marcus, 1973; Martikainen et al., 2005; Aliu et al., 2009; Baess et al., 2011) and temporally predictable sounds which are not self-generated but temporally cued via a visual stimulus (Ford et al., 2007; Sowman et al., 2012; Oestreich et al., 2015).

The underlying functional anatomy engaged during auditory attenuation is yet to be determined. The arcuate fasciculus provides a direct connection between speech production (Broca’s) and speech perception (Wernicke’s) areas and is therefore a plausible white matter connection for the conveyance of efference copies according to internal forward models of auditory attenuation during willed speech. In addition to direct, long segment fibers connecting Broca’s and Wernicke’s area, the arcuate fasciculus also has shorter, indirect connections consisting of an anterior pathway which connects Broca’s area to Geschwind’s territory, and a posterior pathway which connects Geschwind’s territory and Wernicke’s area (Catani et al., 2005). These long and short distance pathways of the arcuate fasciculus possess different functional roles, whereby the direct pathway is thought to be involved in phonologically functions and the indirect pathways in semantic functions (Catani and ffytche, 2005). Specifically, the posterior indirect pathway is thought to be involved in auditory comprehension and the anterior indirect pathway in the vocalization of semantic information (Catani et al., 2005). The arcuate fasciculus has been suggested as the most likely connection to be utilized during auditory attenuation to willed speech (Pynn and DeSouza, 2013).

Additionally, the frontal aslant, which directly connects Broca’s area with the supplementary motor area (Catani et al., 2012), might also play a role in auditory attenuation, as it is involved in verbal fluency (Catani et al., 2013) and speech initiation (Fujii et al., 2016). It is therefore conceivable that these white matter pathways are functionally engaged and effectively connected during auditory attenuation.

In this study, we formulated a set of dynamic causal models (DCMs), which map onto the plausible functional anatomy of auditory attenuation to self-generated and temporally predictable speech. These DCMs included brain regions interconnected via the arcuate fasciculus and the frontal aslant. According to the forward model, efference copies are transmitted via backward connections along the arcuate fasciculus (Pynn and DeSouza, 2013). In keeping with this theory, it was hypothesized that models with both forward and backward connections, which convey sensory input and prediction, respectively, are better at explaining auditory attenuation than models with forward connections alone. Furthermore, we explored whether auditory attenuation was better explained by alternative models that included or excluded the above mentioned regions along the arcuate fasciculus (Geschwind’s territory) and the frontal aslant (supplementary motor area).

## 2. Materials and Methods

### 2.1 Participants

Seventy-five healthy participants (38% males, aged 18-44 years, 95% right-handed) were recruited through the online recruitment systems SONA-1 and SONA-P at the University of New South Wales, Australia. Participants were either monetarily reimbursed for their time or received course credit. One participant was excluded from the analyses due to a self-reported diagnosis of an Axis I disorder (American Psychiatric Association, 2000). Event-related potential (ERP) analyses and a detailed description of the demographic data have been reported previously elsewhere (Oestreich et al., 2015). All participants gave written informed consent. This study was approved by the UNSW Human Research Ethics Advisory Panel (Psychology) and the University of Queensland Research Ethics Committee.

### 2.2 Procedure

Participants completed a number of questionnaires about their demographics, alcohol, nicotine, caffeine and recreational drug use, as well as history of Axis I disorders. Participants then underwent electroencephalographic (EEG) recordings while performing an experimental task in a quiet, dimly lit room. The experiment consisted of three conditions, namely the *Talk*, *Passive Listen*, and *Cued Listen* conditions (Ford et al., 2007; Oestreich et al., 2015). Before the experiment, an instruction video was played, which demonstrated how to vocalize the syllable ‘ah’ in a clear manner while maintaining the gaze on a fixation cross. Following the instruction video, participants were trained to vocalize the syllable ‘ah’ with a duration of less than 300ms and an intensity between 75dB and 85dB. During the *Talk* condition, participants vocalized a series of ‘ah’s in a desk-mounted microphone, every one to three seconds until 3 minutes had elapsed. In the *Cued Listen* condition, participants were instructed to listen to a recording of their own willed vocalizations whilst watching a video of the vocalization waveforms. Participants were therefore able to make exact temporal predictions about the onset of a speech sound. Lastly, during the *Passive Listen* condition, participants listened to their own willed vocalizations played back without a cue. During the *Passive Listen* condition, participants are therefore unable to make temporal predictions about the onset of the next speech sound.

### 2.3 Data Acquisition and preprocessing

EEG was recorded with a 64-channel BioSemi ActiView system at a sampling rate of 2048Hz, 18dB/octave roll-off and 417Hz bandwidth (3dB). External electrodes were placed on the mastoids, the outer canthi of both eyes and below the left eye and the EEG data were referenced to the average of the mastoid electrodes. Preprocessing was performed using SPM12 (Wellcome Trust Centre for Neuroimaging, London; http://www.fil.ion.ucl.ac.uk/spm/) with MATLAB (MathWorks). Triggers were inserted at the onset of each ‘ah’ and the EEG data were then segmented into 800ms intervals with 200ms pre- and 600ms post-stimulus onset. Eye blinks and movements were corrected with a regression based algorithm using vertical and horizontal electrooculogram (VEOG, HEOG; Gratton et al. (1983). The low and high frequency components of the EEG signal were attenuated using a 0.5-15Hz bandpass filter (Ford and Mathalon, 2004) and trials containing artefacts exceeding ±50µV were rejected. The remaining artifact free trials were averaged per condition for each participant in order to obtain event-related potentials (ERPs). ERPs were baseline corrected using the –100-0ms pre-stimulus interval. The N1 component of each ERP was defined as the most negative peak between 50ms and 150ms after the onset of a speech sound.

### 2.4 Source Reconstruction

Source images were obtained by reconstructing scalp activity using a Boundary Element Method (BEM) and a standard MNI template for the cortical mesh, in the absence of individual MRIs. Images from these reconstructions were obtained from each of the three conditions in every participant and smoothed at FWHM 8x8x8 mm^3^. We then performed within-subjects F-test for the main effect of *condition* (Talk/Passive Listen/Cued Listen) and t-tests for the *Talk* vs *Passive Listen* and *Cued Listen* vs *Passive Listen* contrasts over the whole brain 3-dimensional space. Effects are displayed at an uncorrected threshold of p < .001.

### 2.5 Dynamic Causal Modelling (DCM)

DCM relies on a generative spatiotemporal model for EEG responses evoked by experimental stimuli (Kiebel et al., 2008). It uses neural mass models (David and Friston, 2003) to infer source activity of dynamically interacting excitatory and inhibitory neuronal subpopulations (Jansen and Rit, 1995), and the connectivity established amongst different brain regions. DCM sources are interconnected via forward, backward and lateral connections (Felleman and Van Essen, 1991), and are arranged in a hierarchical manner (David et al., 2005; Kiebel et al., 2007). DCMs are designed to test specific connectional hypotheses that are motivated by alternative theories (Garrido et al., 2008). Every connectivity model defines a network that attempts to predict (i.e. generate) the ERP signal. Differences in the ERPs to different experimental stimuli are modelled in terms of synaptic connectivity changes within and between cortical sources (Garrido et al., 2008). Several plausible cortical network connections are compared by estimating the probability of the data given a particular model within the space of models compared, using Bayesian Model Selection (BMS;(Penny et al., 2004). BMS provides estimates of the posterior probability of the DCM parameters given the data, as well as the posterior probability of each model (Penny et al., 2004). The winning model is the model, which maximizes the fit to the data while simultaneously minimizing the complexity of the model.

The posterior probability of each model was computed over all participants using a random effects approach (RFX; (Stephan et al., 2009). The conventional fixed effects approach for model comparison is limited by the assumption that all participants’ data are generated by the same model and is not very robust to outliers. The RFX approach used in the current study on the other hand, is able to quantify the probability that a specific model generated the data for any randomly chosen participant relative to other models. Moreover, RFX is robust to outliers (Stephan et al., 2009). Our main conclusions are based on inferences at the family level with a RFX exceedance probability of .96 on average (ranging from .85-1). In addition to RFX we also report the Bayesian omnibus risk (BOR), which quantifies the risk incurred when performing Bayesian model selection, by directly measuring the probability that all model frequencies are equal (Rigoux et al., 2014). The BOR is a bounded between 0 and 1, whereby a value close to 1 indicates that the models are equal, whereas a value close to 0 indicates that the models are well distinguishable from one another.

### 2.6 Model specification

The models compared in this study include up to 10 brain regions hierarchically organized in one to five levels. These alternative models are motivated by speech related brain regions that are connected via the auditory white matter pathways of the arcuate fasciculus and the frontal aslant, which are thought to be involved in auditory attenuation. Source reconstruction identified these brain regions to be significantly activated during the tasks of this study (see Figure 4). Additionally, previous studies using similar tasks to the present paradigm reported coinciding brain regions to be activated: A study using concurrent EEG and fMRI found the superior temporal gyrus (STG), which includes Wernicke’s area and the primary auditory cortex to be activated (Ford et al., 2016), and a study using EEG with anatomical MRI reported activity in the STG, sensorimotor area and inferior frontal gyrus, which includes Broca’s area (Wang et al., 2014). The bilateral primary auditory cortex (A1) was defined as the cortical input node for auditory sensory information. The arcuate fasciculus consists of a direct pathway between Wernicke’s area (W) and Broca’s area (B) as well as two indirect pathways, namely the posterior pathway connecting W and Geschwind’s territory (G), and the anterior pathway connecting G and B. The frontal aslant interconnects B with the supplementary motor area (SMA). The mean locations of the nodes were based on the Montreal Neurological Institute (MNI) coordinates for left A1 (-52, -19, 7), right A1 (50, -21, 7), left W (-57, -20, 1), right W (54, -19, 1), left G (-53, -32, 33), right G (51, -33, 34), left B (-48, 13, 17), right B (49, 12, 17), left SMA (-28, -2, 52) and right SMA (28, -1, 51; see Figure 1).

**Figure 1.**
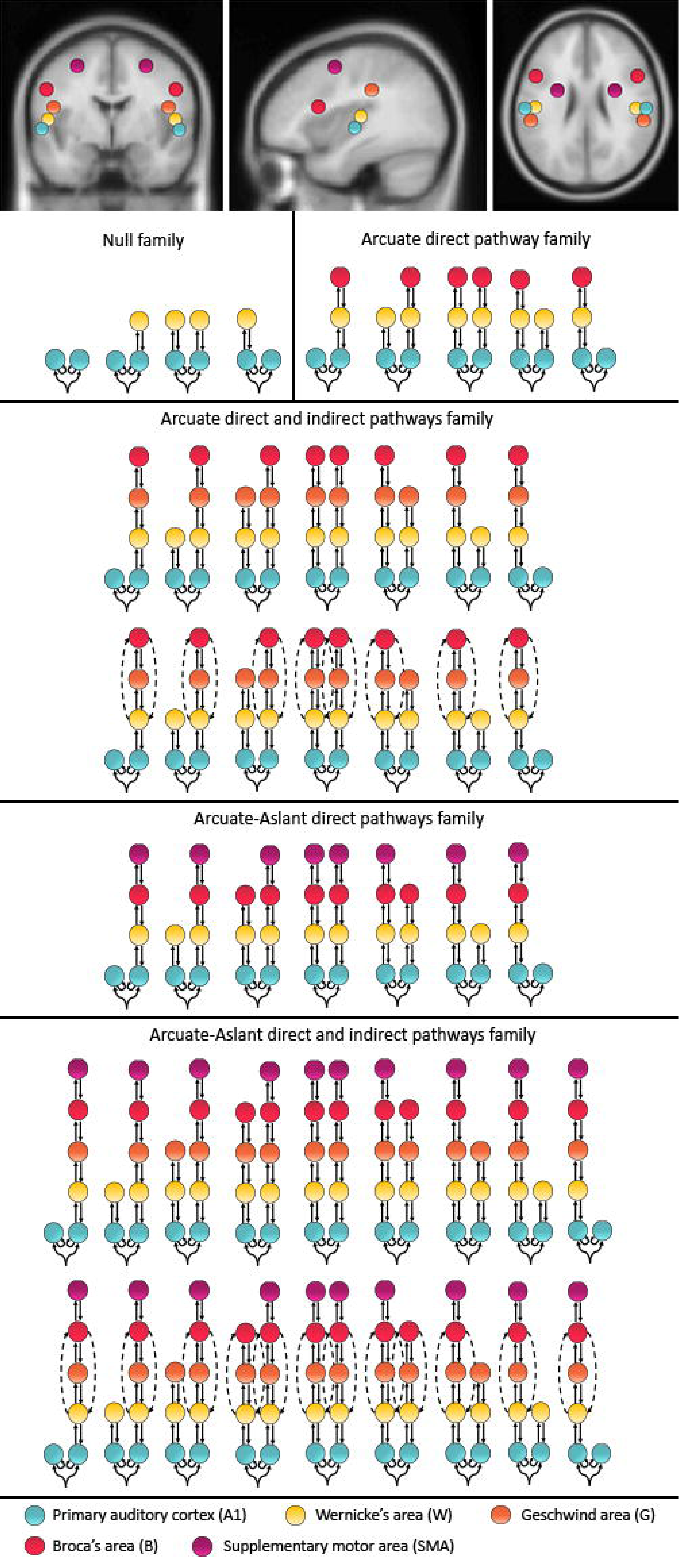
Mean locations for the DCM nodes and model space. The MNI coordinates include: left A1 (-52, -19, 7), right A1 (50, -21, 7), left W (-57, -20, 1), right W (54, -19, 1), left G (-53, -32, 33), right G (51, -33, 34), left B (-48, 13, 17), right B (49, 12, 17), left SMA (-28, -2, 52), SMA (28, -1, 51). The 48 represented models were included twice, once with *Forward* connections only and once with *Forward and backward* connections. These 96 models were chosen to test different hypotheses about the functional anatomy of auditory attenuation to temporally predictable speech. The models were combined to 5 families including a *Null family*, the *Arcuate direct pathway family*, the *Arcuate direct and indirect pathways family*, the *Aslant-Arcuate direct pathways family*, and the *Aslant-Arcuate direct and indirect pathways and Aslant*.

**Figure 2.**
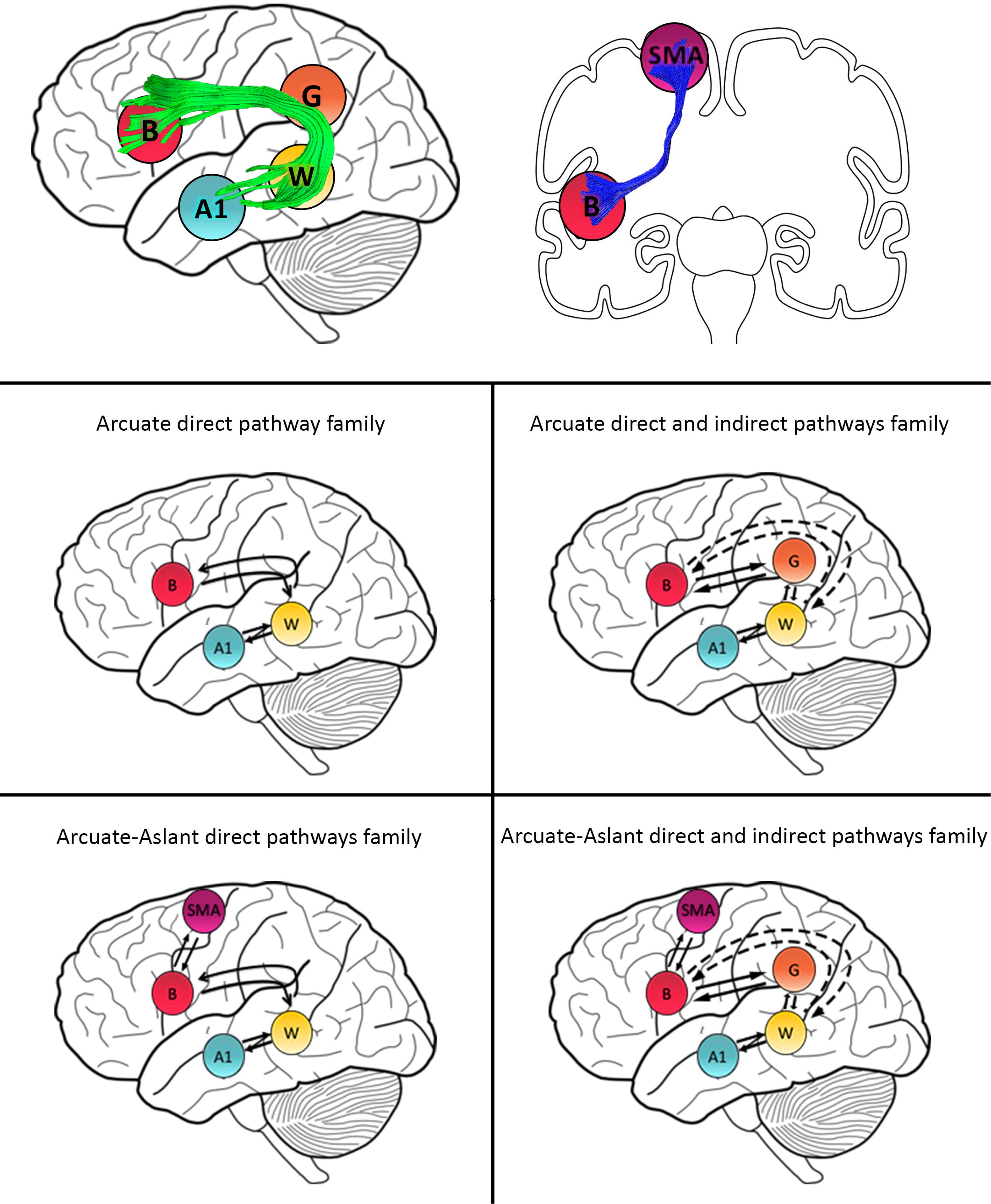
Family definitions and anatomical tracts. Primary auditory cortex (A1), Wernicke’s area (W), Geschwind’s territory (G) and Broca’s area (B) are interconnected via the arcuate fasciculus (green). B and supplementary motor area (SMA) are interconnected by the frontal aslant (blue). Schematic representation of the *Arcuate direct pathway family*, the *Arcuate direct and indirect pathways family*, the *Aslant-Arcuate direct pathways family*, and the *Aslant-Arcuate direct and indirect pathways and Aslant*.

**Figure 3.**
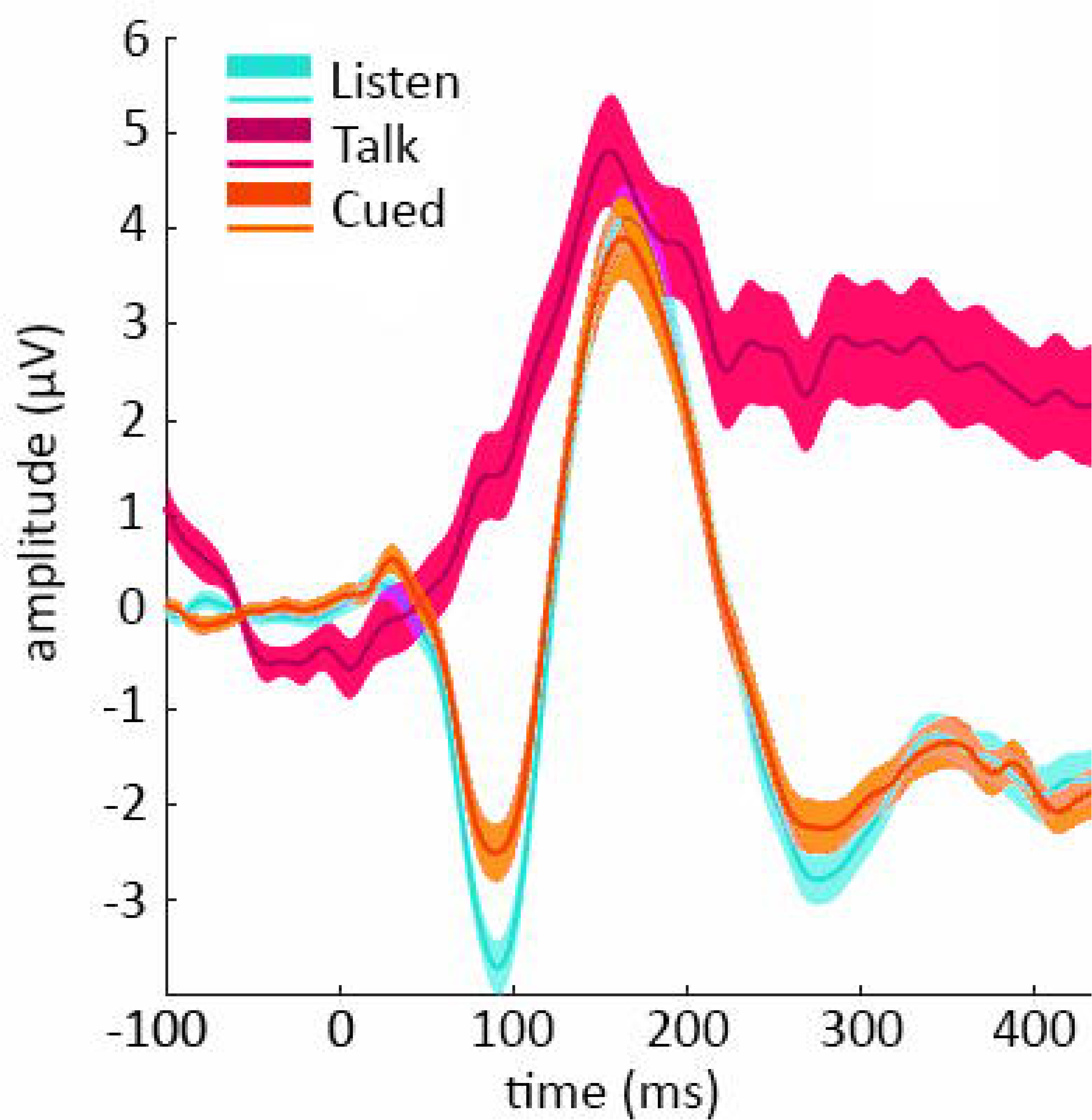
Event-related potentials (ERPs) from electrode Cz in response to willed vocalization in the Talk (magenta), Cued Listen (orange) and Passive Listen (cyan) conditions.

**Figure 4.**
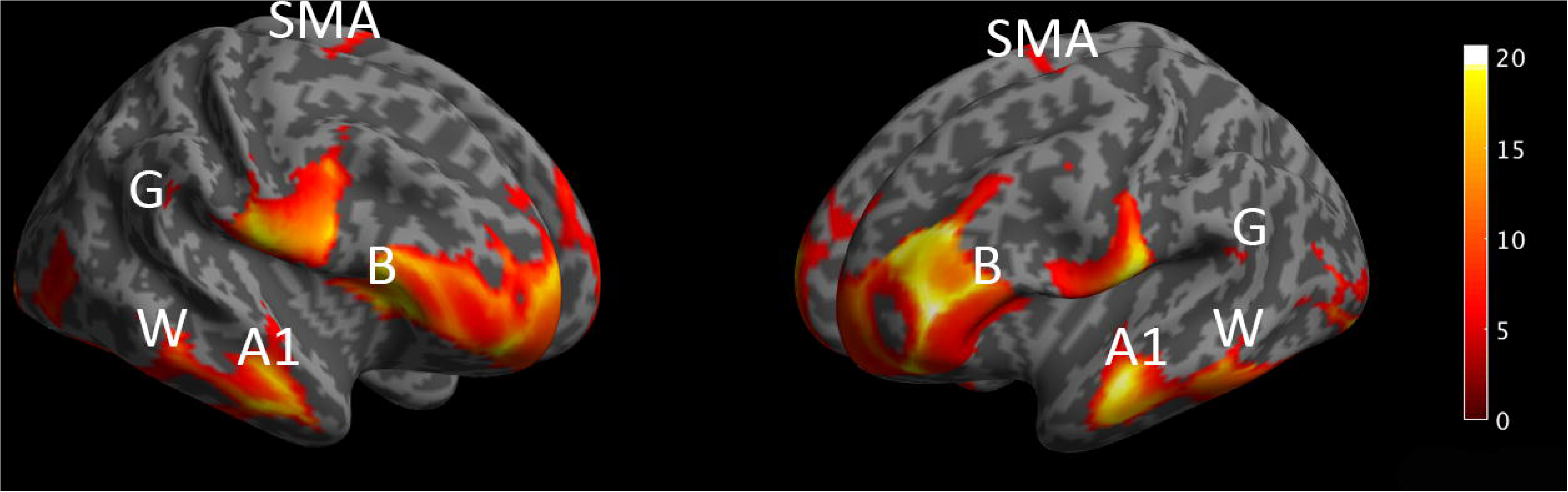
Source reconstruction analysis revealed several significant clusters for the main effect of *condition* (Talk/Passive Listen/Cued Listen) at p < .001, uncorrected. Significant clusters include bilateral primary auditory cortex (A1), Wernicke’s area (W), Geschwind’s territory (G) Broca’s area (B) and supplementary motor area (SMA).

Since the effective connectivity of auditory attenuation has not been studied before, we considered a comprehensive model space including a total of 96 models comprising symmetric and non-symmetric hierarchical models, with forward connections only and combined forward and backward connections, with and without indirect connections between W and B via G, as well as models with and without the frontal aslant, which connects B to SMA (for a full description of the model space see Figure 1). All models allowed for changes of intrinsic connectivity at the level of A1. All 96 models were estimated and individually compared to each other using BMS. The 96 models were then partitioned into a number of different families.

We investigated whether auditory attenuation is driven by feedback loops, through both forward and backward connections, or by bottom-up inputs alone, via forward connections between brain regions along the arcuate fasciculus, and possibly also through the frontal aslant. Models with feedback loops would support the theory of internal forward models whereby self-generated actions and predictable sensory consequences are cortically attenuated. To this end, a family consisting of all 48 models with *Forward family* connections only was compared to a family consisting of all 48 models with *Forward and Backward family* connections.

We then grouped our models into families that included specific regions and tracts as follows: 1) The *Null family* consisted of 8 models that included A1 only and models connecting A1 to W, 2) the *Arcuate direct pathway family* included 10 models, with connections between A1 and W as well as W and B, 3) the *Arcuate direct and indirect pathways family* consisted of 28 models including connections between A1 and W, W and G, G and B, as well as W and B, 4) The *Aslant-Arcuate direct pathways family* included 14 models that connected A1 and W, W and B, as well as B and SMA and 5) the *Aslant-Arcuate direct and indirect pathways and Aslant* comprising 18 models, included connections between A1 and W, W and G, G and B, W and B as well as B and SMA (see Figure 1 and 2).

To follow up whether models with or without the frontal aslant (i.e. connections to SMA) better explained auditory attenuation, we first combined the *Arcuate direct pathway family* (10 models with connections linking A1, W, and B directly; see Figure 1 and 2) and the *Arcuate direct and indirect pathways family* (28 models linking A1, W, G and B) into one single family – the *Arcuate family.* We then compared this to the *Arcuate-Aslant family*, which resulted from combining the *Arcuate-Aslant direct pathways family* (14 models) and the *Arcuate-Aslant direct and indirect pathways families* (36 models) consisting of all the 50 models with connections to SMA (see Figure 1 and 2).

Lastly, to investigate whether Geschwind’s territory is part of the circuit engaged in auditory attenuation of speech, we compared families of models with and without Geschwind. To this end, we combined all the models precluding Geschwind into one family – *no Geschwind family –* by grouping the *Arcuate direct pathway family* (10 models) and the *Arcuate-Aslant direct pathways family* (14 models; see Figure 1 and 2). We compared the *no Geschwind family* to the *Geschwind family*, which included a combination of the *Arcuate direct and indirect pathways family* (28 models) and the *Arcuate-Aslant direct and indirect pathways family*, that is, all the models that included Geschwind (36 models).

Each of the 96 models was fitted to each individual participant’s mean response for the contrast between the *Passive Listen* and *Talk* conditions (i.e. effective connectivity of self-generated speech), as well as to the contrast between the *Passive Listen* and *Cued Listen* conditions (i.e. effective connectivity of predictable speech), whereby the *Passive Listen* condition was used as the baseline condition for both DCM contrasts.

## 3. Results

### 3.1 Scalp analysis

A repeated measures analysis of variances (ANOVA) detected a main effect of *condition* (Talk/Passive Listen/Cued Listen) on the N1-amplitude at electrode Cz [*F*(2,144) = 20.297, *p* = .000002, 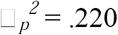] (see Figure 3). Bonferroni-corrected follow-up contrasts revealed a significant difference between the Talk and the Passive Listen conditions [*F*(1,72) = 26.969, *p* = .000004, 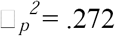] and the Cued and Passive Listen conditions [*F*(1,72) = 6.051, *p* = .032,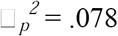]. A detailed analyses of the event-related potentials (ERP) has been reported previously elsewhere (Oestreich et al., 2015).

### 3.2 Source Reconstruction

We observed a main effect of *condition* (Talk/Passive Listen/Cued Listen at *p* < .001, uncorrected) in the bilateral inferior frontal gyrus (which includes Broca’s area, B), superior temporal gyrus (comprising the primary auditory cortex, A1, and Wernicke’s area, W), medial frontal cortex (including the supplementary motor area, SMA) and the inferior parietal lobule (which is also known as Geschwind’s territory, G; see Figure 4).

### 3.3 DCM analyses

In a first step all 96 models with forward connections only as well as forward and backward connections were individually compared to each other. The DCM analysis of the self-generated speech condition (compared to the *Passive Listen* condition) indicated that the best model included reciprocal connections between bilateral A1, W, G, B and SMA, as well as direct connections between W and B in both hemispheres (expected probability = .04, exceedance probability = .21; BOR = .01, see Figure 5). This indicates bilateral connectivity along the arcuate fasciculus and the frontal aslant best explain attenuation of self-generated speech. The second best model, which was also rather probable, was equal to the winning model except that it did not include any connections to SMA via the aslant (expected probability = .03, exceedance probability = .17; see Figure 5). The DCM analysis of the predictable speech condition (compared to the *Passive Listen* condition) revealed that the winning model was the same as the second most likely model for self-generated speech, including reciprocal connections linking bilateral A1, W, G and B, as well as direct connections between W and B in both the left and the right hemispheres (expected probability = .04, exceedance probability = .32; BOR < .01, see Figure 5). The second best model for predictable speech was in all equal to the winning model except that it included connections to SMA via the aslant in the left hemisphere (expected probability = .03, exceedance probability = .17; see Figure 5). The very low values of the BOR for both model comparisons indicates that the models are well distinguishable from one another, which adds to the confidence in the winning models.

**Figure 5.**
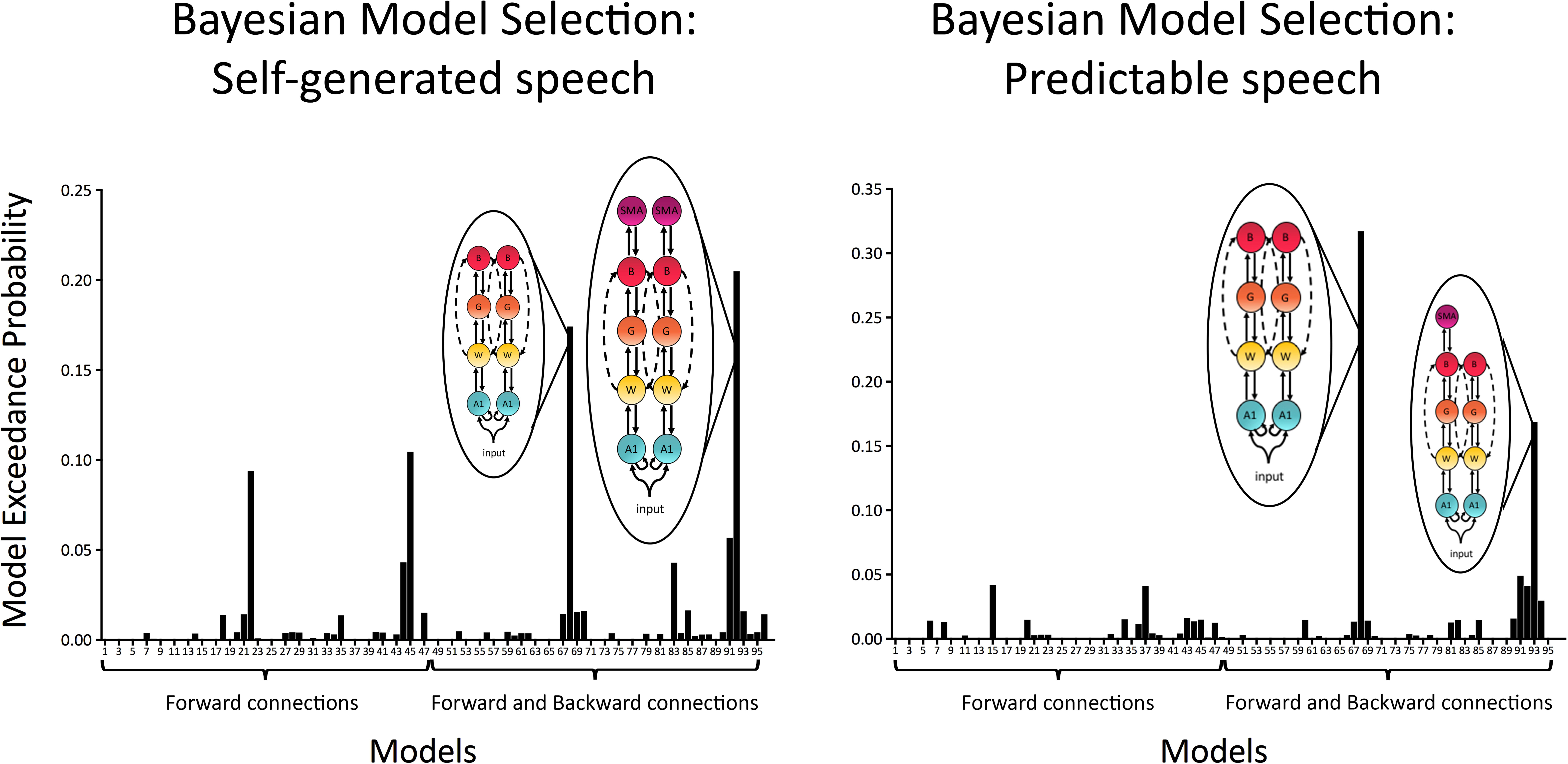
Model exceedance probability for attenuation of self-generated and predictable speech. Bayesian model selection (Random effects) over the whole model space indicated self-generated speech attenuation was best explained by a model with recurrent (i.e., forward and backward) modulations between bilateral primary auditory cortex (A1), Wernicke’s area (W), Geschwind’s territory (G) Broca’s area (B) and supplementary motor area (SMA), as well as direct bilateral connections between W and B. This model was followed by a model, which was in all equal to the winning model except that it did not include bilateral connections from B to SMA. Attenuation of predictable speech was best explained by a model with recurrent connections between bilateral A1, W, G and B, as well as direct bilateral connections between W and B. This model was followed by a model, which was in all equal to the winning model except that it included a connection from B to SMA in the left hemisphere.

When comparing a family with modulations of forward connections only (i.e. *Forward family*) to a family of both forward and backward connections (i.e. *Forward and Backward family*), we found that the family consisting of a combination of *Forward and Backward* connections better explained auditory attenuation for both self-generated speech (expected probability = .59, exceedance probability = .95) and temporally predictable speech (expected probability = .56, exceedance probability = .85), than the families including *Forward* connections only (see Figure 6).

**Figure 6.**
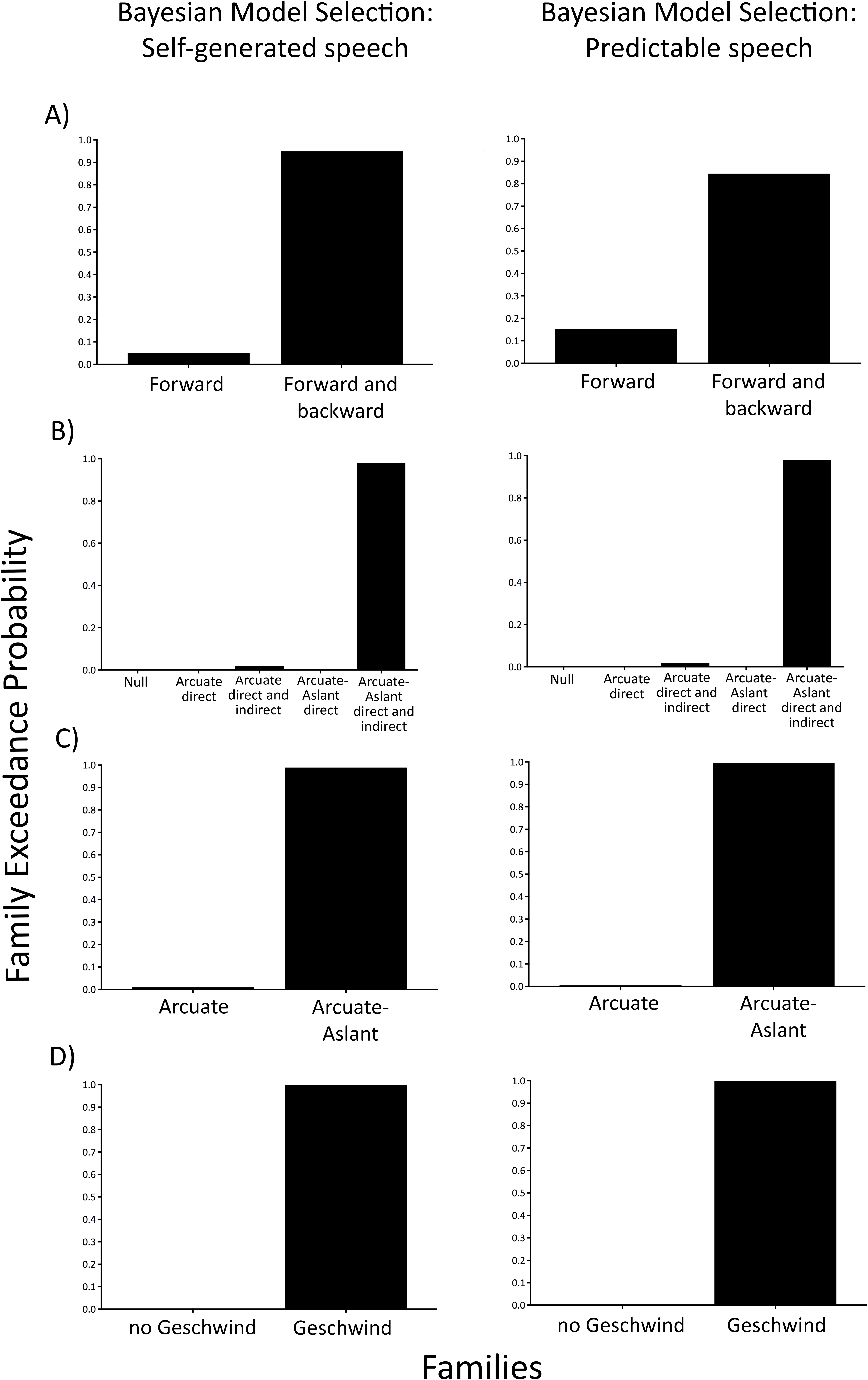
Family-level inference for attenuation of self-generated and predictable speech sounds –exceedance probabilities for the family comparisons. A) comparison of the *Forward family* (48 models) to the *Forward and Backward family* (48 models). B) comparison of 5 families including a *Null family* (8 models), the *Arcuate direct pathway family* (10 models), the *Arcuate direct and indirect pathways family* (28 models), the *Aslant-Arcuate direct pathways family* (14 models), and the *Aslant-Arcuate direct and indirect pathways and Aslant* (18 models). C) comparison of the *Arcuate family* (38 models) and the *Arcuate-Aslant family* (50 models). D) comparison of *no Geschwind family* (24 models) to *Geschwind family* (64 models).

To test specific hypotheses as to which brain regions interconnected by the arcuate fasciculus and the frontal aslant were engaged in auditory attenuation, five families of models were created as described in the methods section (see Figure 1 and 2). BMS of these families indicated that the *Aslant-Arcuate direct and indirect pathways family* was the winning family for both self-generated speech (expected probability = .55, exceedance probability = .98) and predictable speech (expected probability = .54, exceedance probability = .98; see Figure 6).

When comparing families with the arcuate fasciculus alone (i.e *Arcuate family*) to families including both the arcuate fasciculus and the frontal aslant (i.e. *Arcuate-Aslant family*), BMS revealed that the winning family *Arcuate-Aslant family* was much more likely than the *Arcuate family* during self-generated speech (expected probability = .63, exceedance probability = .99) and predictable speech (expected probability = .60, exceedance probability = .95; see Figure 6).

Lastly, we investigated whether families of models with or without Geschwind, which enquired as to whether Geschwind plays a role in the functional circuit engaged in auditory attenuation (*Geschwind family* vs *no Geschwind family*). Our results indicated that the family of models including connections to Geschwind outperformed models without Geschwind during self-generated speech (expected probability = .91, exceedance probability = 1) and predictable speech (expected probability = .88, exceedance probability = 1; see Figure 6).

## 4. Discussion

This study investigated the functional anatomy underlying auditory attenuation to self-generated and temporally predictable speech sounds using DCM. Model comparison revealed that modulations in both forward and backward connections better explained auditory attenuation than forward connections alone, which is in line with the theory of internal forward models of auditory attenuation (Ford and Mathalon, 2004), whereby an efference copy, or prediction, is conveyed via backward (i.e. top-down) connections. Connectivity models linking primary auditory cortex, Wernicke’s area, Geschwind’s territory and Broca’s area via the arcuate fasciculus and the supplementary motor area, through the frontal aslant tract, outperformed models without connections to the supplementary motor area and Geschwind’s territory. These findings indicate that the circuitry underlying auditory attenuation to self-generated and temporally predictable speech sounds most likely involves brain regions interconnected by both the short distance, indirect pathways, and the long distance, direct pathway of the arcuate fasciculus in addition to brain regions interconnected by the frontal aslant.

The finding that a combination of forward and backward connections better explained auditory attenuation than forward connections alone is in line with the theory of internal forward models, whereby a prediction, in the form of an efference copy, is conveyed through backward connections. Forward connections can be conceptualized as bottom-up processes (Friston, 2005; Chen et al., 2009), which convey environmental sensory information from the primary auditory cortex to higher cortical levels. On the contrary, backward connections represent top-down (Chen et al., 2009), predictive processes based on self-monitoring or past experiences. In this study, we used a *Talk* condition during which participants vocalized speech sounds and a *Cued Listen* condition whereby participants were cued to the exact onset of each speech sound while listening to the previously recorded vocalizations from the *Talk* condition. We compared these conditions to the baseline, *Passive Listen* condition, during which participants were passively listening to the series of previously recorded vocalizations from the *Talk* condition. During the *Talk* and *Cued Listen* conditions, participants are able to make predictions about each speech sound, via top-down, backward connections, which are then compared with actual sensory inputs sent upwards via forward connections. On the contrary, during the *Passive Listen* condition participants are unable to make temporal predictions about to the onset of a speech sound. The theory of internal forward models is therefore supported by the findings from this study, whereby changes in effective connectivity from the baseline (i.e., *Passive Listen* condition) to the *Talk* conditions is best explained by a feedback loop comprising conjoint forward (bottom-up) and backward (top-down) connections. Similarly, feedback loops were found to underlie connectivity differences between the *Cued Listen* compared to the *Passive Listen* condition, suggesting that a forward model for predictable sounds is also internally generated and conveyed via backward connections.

The arcuate fasciculus has been proposed as the most likely route for the efference copy of a motor act during internal forward models of auditory attenuation (Whitford et al., 2011; Pynn and DeSouza, 2013). On the contrary, the frontal aslant seems to be a likely connection for the initiation of the motor act to trigger willed speech as it has a connection to the supplementary motor area. The results from the individual model comparisons support this theory, as the winning model during self-generated speech included bilateral connections along the arcuate fasciculus and the frontal aslant, thereby facilitating the transmission of an efference copy and a motor efference. During temporally predictable speech on the other hand, the most probable model included connections along the arcuate fasciculus only.

In order to determine whether the frontal aslant adds to the functional anatomy of auditory attenuation or whether connections along the arcuate fasciculus alone are sufficient, we compared families of all models with and without connections along the frontal aslant (while keeping the arcuate fasciculus pathways intact). The findings indicated that during both self-generated speech and predictable speech, models with connections along the arcuate fasciculus and the frontal aslant better explained auditory attenuation than models including the arcuate fasciculus only. This can be explained by the motor efference in the *Talk* condition, whereby speech sounds are actively generated and the motor efference is sent along the frontal aslant. However, the *Cued Listen* condition did not involve a motor act, which means that the frontal aslant tract is not being utilized for the transmission of a motor efference. A possible explanation for the involvement of connections to the supplementary motor area during predictable speech and therefore the engagement of the frontal aslant is the notion of an efference copy for thought. Indeed, a proposal put forward by (Jackson, 1958) states that since internal forward models are working reliably during processes of sensory motor control, the same internal forward models, developed later in evolution, might also be utilized during higher cognitive processes such as thought or inner speech, which can be seen as our most complex motor act without actions. In the context of the present study, while participants are not actively generating the vocalization in the *Cued Listen* condition, watching the waveforms of the speech sounds might lead them to internally simulate the next vocalization, which might explain the activation of the supplementary motor area without a motor act.

The arcuate fasciculus consists of long distance fibers which connect Broca’s and Wernicke’s area as well as short distance fibers which connect Broca’s and Geschwind’s territory via an anterior pathway, and Geschwind’s territory and Wernicke’s area via a posterior pathway (Catani et al., 2005). The results of the present study indicate that models including long distance connections in addition to short distance connections, via Geschwind’s territory, better explain auditory attenuation than models including long distance connections only. The direct, long distance pathway is thought to be involved in phonological repetitions (Catani and ffytche, 2005) and therefore represents a plausible connection to be utilized during this experimental tasks, whereby the same speech sound was vocalized and played repetitively. The indirect, short distance pathways of the arcuate fasciculus are thought to be involved in semantic functions (Catani and ffytche, 2005). The engagement of these connections during auditory attenuation might be explained by the nature of the speech sounds used in the present study. Since phonemes are the building blocks of language which are used to distinguish one word from another, it is possible that participants assigned semantic meaning to these sounds, which would likely not occur if the sounds were simple tones.

The involvement of brain areas interconnected via the arcuate fasciculus during auditory attenuation is in line with findings from studies of auditory attenuation in schizophrenia. There is substantial evidence that patients with schizophrenia possess abnormal auditory attenuation to self-generated speech (Ford et al., 2001; Ford and Mathalon, 2004; Ford et al., 2007), button-press elicited sounds (Whitford et al., 2011; Ford et al., 2014), and temporally cued sounds (Ford et al., 2007). Individuals at high-risk for developing a psychotic disorder exhibit auditory attenuation that is intermediate between healthy participants and patients with schizophrenia (Perez et al., 2012). Moreover, healthy individuals with psychotic-like experiences show less auditory attenuation compared to healthy individuals without psychotic-like experiences (Oestreich et al., 2015, 2016).

The underlying mechanisms inducing these auditory attenuation deficits in schizophrenia and psychosis are still unclear. However, several studies have reported changes to the white matter structure and specifically to the myelin sheath of the axons constituting the arcuate fasciculus in patients with schizophrenia. This is insofar important as it indicates that connectivity along the arcuate fasciculus during auditory attenuation should be delayed due to a loss of conduction velocity induced by demyelination. Support for this contention comes from a study by (Whitford et al., 2011), which reported that auditory attenuation abnormalities typically exhibited by patients with schizophrenia could be completely eliminated by imposing a 50ms delay between a self-generated button press and the delivery of a sound. This was interpreted to indicate that efference copies travelling along the arcuate fasciculus during auditory attenuation were delayed by 50ms in the group of schizophrenia patients. Furthermore, the study reported that the degree to which auditory attenuation improved as a result of the delay between button press and tone delivery was linearly correlated with white matter abnormalities in the arcuate fasciculus. The findings from the present study add further support for the role of the arcuate fasciculus during auditory attenuation by showing that the brain regions that are structurally interconnected by the arcuate fasciculus are also effectively connected.

In summary, our study shows that auditory attenuation for self-generated and predictable speech involve brain regions such as Wernicke’s area, Broca’s area, and Geschwind’s territory, interconnected through the arcuate fasciculus via both short and long distance fibers, as well as the supplementary motor area, which is linked to Broca’s area via the frontal aslant. Critically, we found that auditory attenuation to self-generated and temporally predictable speech sounds engages feedback loops with conjoint forward (bottom-up) and backward (top-down) connections. This is consistent with internal forward models, whereby the sensory consequences of our actions and thoughts are internally predicted, which is reflected in reduced cortical responsiveness to these sensations.

## References

Aliu SO, Houde JF, Nagarajan SS (2009) Motor-induced suppression of the auditory cortex. Journal of Cognitive Neuroscience 21:791–802.

American Psychiatric Association (2000) Diagnostic and statistical manual of mental disorders. Washington, DC: Author.

Baess P, Horváth J, Jacobsen T, Schröger E (2011) Selective suppression of self-initiated sounds in an auditory stream: An ERP study. Psychophysiology 48:1276–1283.

Blakemore S, Wolpert DM, Frith CD (1998) Central cancellation of self-produced tickle sensation. Nature Neuroscience 1:635–640.

Catani M, ffytche DH (2005) The rises and falls of disconnection syndromes. Brain 128:2224–2239.

Catani M, Jones DK, ffytche DH (2005) Perisylvian language networks of the human brain. Ann Neurol 57:8–16.

Catani M, Dell’acqua F, Vergani F, Malik F, Hodge H, Roy P, Valabregue R, Thiebaut de Schotten M (2012) Short frontal lobe connections of the human brain. Cortex 48:273–291.

Catani M, Mesulam MM, Jakobsen E, Malik F, Martersteck A, Wieneke C, Thompson C, Thiebaut de Schotten M, Dell’ Acqua F, Weintraub S, Rogalski E (2013) A novel frontal pathway underlies verbal fluency in primary progressive aphasia. Brain 136:2619–2628.

Chen CC, Henson RN, Stephan KE, Kilner JM, Friston KJ (2009) Forward and backward connections in the brain: a DCM study of functional asymmetries. Neuroimage 45:453–462.

Creutzfeldt O, Ojernan G, Lettich E (1989) Neuronal activity in the human lateral temporal lobe II. Responses to the subject’s own voice. Experimental Brain Research 77:476–489.

Curio G, Neuloh G, Numminen J, Jousmäki V, Hari R (2000) Speaking modifies voice-evoked activity in the human auditory cortex. Hum Brain Mapp 9:183–191.

David O, Friston KJ (2003) A neural mass model for MEG/EEG: coupling and neuronal dynamics. Neuroimage 20:1743–1755.

David O, Harrison L, Friston KJ (2005) Modelling event-related responses in the brain. Neuroimage 25:756–770.

Eliades SJ, Wang X (2003) Sensory-motor interaction in the primate auditory cortex during self-initiated vocalizations. J Neurophysiol 89:2194–2207.

Feinberg I (1978) Efference copy and corollary discharge - implications for thinking and its disorders. Schizophr Bull 4:636–640.

Felleman DJ, Van Essen DC (1991) Distributed hierarchical processing in the primate cerebral cortex. Cerebral Cortex 1:1–47.

Ford JM, Mathalon DH (2004) Electrophysiological evidence of corollary discharge dysfunction in schizophrenia during talking and thinking. J Psychiatr Res 38:37–46.

Ford JM, Palzes VA, Roach BJ, Mathalon DH (2014) Did I do that? Abnormal predictive processes in schizophrenia when button pressing to deliver a tone. Schizophr Bull 40:804–812.

Ford JM, Roach BJ, Palzes VA, Mathalon DH (2016) Using concurrent EEG and fMRI to probe the state of the brain in schizophrenia. Neuroimage Clinical 12:429–441.

Ford JM, Gray M, Faustman WO, Roach BJ, Mathalon DH (2007) Dissecting corollary discharge dysfunction in schizophrenia. Psychophysiology 44:522–529.

Ford JM, Mathalon DH, Heinks T, Kalba S, Faustman WO, Roth WT (2001) Neurophysiological Evidence of Corollary Discharge Dysfunction in Schizophrenia. Am J Psychiatry 158:2069–2071.

Friston KJ (2005) A theory of cortical responses. Philosophical transactions of the Royal Society of London 360.

Fujii M, Maesawa S, Ishiai S, Iwami K, Futamura M, Saito K (2016) Neural Basis of Language: An Overview of An Evolving Model. Neurol Med Chir 56:379–386.

Garrido MI, Friston KJ, Kiebel SJ, Stephan KE, Baldeweg T, Kilner JM (2008) The functional anatomy of the MMN: a DCM study of the roving paradigm. NeuroImage 42:936–944.

Gratton G, Coles MG, Donchin E (1983) A new method for off-line removal of ocular artifact. Electroencephalography and Clinical Neurophysiology 55:468–484.

Heinks-Maldonado TH, Mathalon DH, Gray M, Ford JM (2005) Fine-tuning of auditory cortex during speech production. Psychophysiology 42:180–190.

Jackson JH (1958) Selected writings of John Hughlings Jackson. New York, NY: Basic Books.

Jansen BH, Rit VG (1995) Electroencephalogram and visual evoked potential generation in a mathematical model of coupled cortical columns. Biological Cybernetics 73:357–366.

Kiebel SJ, Garrido MI, Friston KJ (2007) Dynamic causal modelling of evoked responses: the role of intrinsic connections. Neuroimage 36:332–345.

Kiebel SJ, Garrido MI, Moran RJ, Friston KJ (2008) Dynamic causal modelling for EEG and MEG. Cognitive Neurodynamics 2:121–136.

Martikainen MH, Kaneko K, Hari R (2005) Suppressed responses to self-triggered sounds in the human auditory cortex. Cerebral Cortex 15:199–302.

Oestreich LKL, Mifsud NG, Ford JM, Roach BJ, Mathalon DH, Whitford TJ (2015) Subnormal sensory attenuation to self-generated speech in schizotypy: Electrophysiological evidence for a ’continuum of psychosis’. Int J Psychophysiol 97:131–138.

Oestreich LKL, Mifsud NG, Ford JM, Roach BJ, Mathalon DH, Whitford TJ (2016) Cortical suppression to delayed self-initiated auditory stimuli in schizotypy: neurophysiological evidence for a continuum of psychosis. Clin EEG Neurosci 47:3–10.

Penny WD, Stephan KE, Mechelli A, Friston KJ (2004) Comparing dynamic causal models. Neuroimage 22:1157–1172.

Perez VB, Ford JM, Roach BJ, Loewy RL, Stuart BK, Vinogradov S, Mathalon DH (2012) Auditory cortex responsiveness during talking and listening: early illness schizophrenia and patients at clinical high-risk for psychosis. Schizophr Bull 38:1216–1224.

Pynn L, DeSouza J (2013) The function of efference copy signals: implications for symptoms of schizophrenia. . Vision Research 76:124–133.

Rigoux L, Stephan KE, Friston KJ, Daunizeau J (2014) Bayesian model selection for group studies - revisite. Neuroimage 84:971–985.

Schafer EW, Marcus MM (1973) Self-stimulation alters human sensory brain responses. Science 181:175–177.

Sowman PF, Kuusik A, Johnson BW (2012) Self-initiation and temporal cueing of monaural tones reduce the auditory N1 and P2. Experimental Brain Research 222:149–157.

Sperry RW (1950) Neural basis of the spontaneous optokinetic response produced by visual inversion. Journal of Comparative and Physiological Psychology 43:482–489.

Stephan KE, Penny WD, Daunizeau J, Moran RJ, Friston KJ (2009) Bayesian model selection for group studies. Neuroimage 46:1004–1017.

Wang J, Mathalon DH, Roach BJ, Reilly J, Keedy SK, Sweeney JA, Ford JM (2014) Action planning and predictive coding when speaking. Neuroimage 91:91–98.

Whitford TJ, Mathalon DH, Shenton ME, Roach BJ, Bammer R, Adcock RA, Bouix S, Kubicki M, De Siebenthal J, Rausch AC, Schneiderman JS, Ford JM (2011) Electrophysiological and diffusion tensor imaging evidence of delayed corollary discharges in patients with schizophrenia. Psychol Med 41:959–969.

Wolpert DM, Ghahramani Z, Jordan MI (1995) An internal model for sensorimotor integration. Science 269:1880–1882.

Zouridakis G, Simos PG, Papanicolaou AC (1998) Multiple bilaterally asymmetric cortical sources account for the auditory N1m component. Brain Topogr 10:183–189.

